# Recombinant SARS-CoV-2 lacking initiating and internal methionine codons within ORF10 is attenuated *in vivo*

**DOI:** 10.1101/2023.08.04.551973

**Authors:** Shichun Gu, Eleanor G Bentley, Rachel I Milligan, Abdulaziz M. Almuqrin, Parul Sharma, Adam Kirby, Daniele F Mega, Anja Kipar, Max Erdmann, James Bazire, Kate J. Heesom, Philip A Lewis, I’ah Donovan-Banfield, Charlotte Reston, Isobel Webb, Simon De Neck, Xaiofeng Dong, Julian A Hiscox, Andrew D Davidson, James P Stewart, David A. Matthews

## Abstract

SARS-CoV-2 has been proposed to encode ORF10 as the 3’ terminal gene in the viral genome. However, the potential role and even existence of a functional ORF10 product has been the subject of debate. There are significant structural features in the viral genomic RNA that could, by themselves, explain the retention of the ORF10 nucleotide sequences without the need for a functional protein product. To explore this question further we made two recombinant viruses, firstly a control virus (WT) based on the genome sequence of the original Wuhan isolate and with the inclusion of the early D614G mutation in the Spike protein. We also made a second virus, identical to WT except for two additional changes that replaced the initiating ORF10 start codon and an internal methionine codon for stop codons (ORF10KO). Here we show that the two viruses have apparently identical growth kinetics in a VeroE6 cell line that over expresses TMPRSS2 (VTN cells). However, in A549 cells over expressing ACE2 and TMPRSS2 (A549-AT cells) the ORF10KO virus appears to have a small growth rate advantage. Growth competition experiments were used whereby the two viruses were mixed, passaged in either VTN or A549-AT cells and the resulting output virus was sequenced. We found that in VTN cells the WT virus quickly dominated whereas in the A549-AT cells the ORF10KO virus dominated. We then used a hamster model of SARS-CoV-2 infection and determined that the ORF10KO virus has attenuated pathogenicity (as measured by weight loss). We found an almost 10-fold reduction in viral titre in the lower respiratory tract for ORF10KO vs WT. In contrast, the WT and ORF10KO viruses had similar titres in the upper respiratory tract. Sequencing of viral RNA in the lungs of hamsters infected with ORF10KO virus revealed that this virus frequently reverts to WT. Our data suggests that the retention of a functional ORF10 sequence is highly desirable for SARS-CoV-2 infection of hamsters and affects the virus’s ability to propagate in the lower respiratory tract.

## Introduction

The first manuscript to describe the genome sequence of SARS-CoV-2 identified the major open reading frames (ORFs) of the virus and alongside genes with clear homology to related coronaviruses, a novel ORF10 was described (1). This putative ORF10 is located 3’ proximal to the N coding region and encodes a peptide 38 amino acids long which has no homology with any previously reported protein. Since the initial description of ORF10, there have been a number of manuscripts with conflicting conclusions on whether there is a specific ORF10 mRNA, whether the ORF10 protein is actually made and, if so, is it functionally relevant (2, 3, 4, 5, 6, 7). Analysis of the evolution of the genome sequence in this region has yielded arguments both for (8) and against (6) ORF10 being expressed as a functional protein.

Initially we and others noted that evidence for a specific ORF10 mRNA was rare or undetectable (2, 3, 9). However, an in-depth analysis of the SARS-CoV-2 transcriptome using a novel mRNA re-capping technique coupled with direct RNA sequencing did substantiate the existence of dedicated ORF10 mRNA (10). Others have sought to understand the function of the proposed ORF10 protein through interactome based approaches (11) and suggested that overexpressed ORF10 protein interacts with Cullin-2-RING-ligase complex but this interaction was later shown to apparently have no known effect or function (7, 12). Other teams have pointed out that SARS-CoV-2 variants with premature stop codons within ORF10 do naturally occur and that these variants have no discernible effect on pathogenesis in humans and did not affect viral growth in VeroE6 cells in the laboratory compared to the wild type virus (5). Conversely, it has also been shown that over expression of ORF10 protein leads to mitochondrial localisation of the ORF10 protein where it affects mitochondrial activated viral signalling protein (MAVS) to supress the innate immune response (13). In addition, the ORF10 protein has been proposed to antagonise STING mediated induction of interferon (14) and affect cilia function (12).

Nonetheless, the overwhelming majority of SARS-CoV-2 isolates appear to have an intact ORF10 sequence and ORF8 is more likely to be lost from an isolate than ORF10 (5). Another possibility is that the nucleotide sequence in the region covered by ORF10 is conserved because it plays a role in RNA folding. This region of the viral genome is highly structured and contains folded RNA structures known to contribute to viral innate immune evasion in other viral systems (15, 16, 17, 18, 19, 20, 21, 22). Most recently it has been proposed that ribosome binding to an internal AUG within the ORF10 gene sequence is key to this region forming a functional regulon structure in the RNA that recruits EPRS1 which in turn modulates ribosomal frameshifting in pp1ab (23).

However, to date most of the data on a possible function for the ORF10 protein derives from deliberate overexpression in cell culture which could produce artefactual effects, especially given the likely rarity of the ORF10 mRNA and the lack of direct evidence for ORF10 protein expression in the context of viral infection. Indeed, the recent evidence of a possible regulon within the 3’UTR of the SARS-CoV-2 genome that requires ribosome binding within ORF10 is also based on the analysis of *in vitro* transcripts and not on modified SARS-CoV-2 virus. To shed further light on whether the ORF10 protein or RNA structure (or both) is functionally required in the context of viral replication, we constructed a recombinant virus that eliminated the possibility of ORF10 protein expression. We initially made a control recombinant virus with a Wuhan like background but also containing the D614G mutation in the spike protein (WT). We then used this WT backbone to make a recombinant virus with the initiating AUG changed to UAA and the internal AUG changed to UAG, referred to as ORF10KO. That both WT and ORF10KO viruses were made by reverse genetics techniques gives greater certainty about the genetic diversity of the initial viral stocks.

We examined the proteomic effect of infection with the WT virus compared to ORF10KO and we investigated the growth rates of the two viruses in different cell lines before performing competition assays in two different cell lines. We found that indeed the ORF10KO virus has distinct growth properties in different cell lines – the ORF10KO virus has a growth advantage in A549 cells over expressing human ACE2 and TMPRSS2 (A549-AT cells) but conversely is hampered in VeroE6 cells overexpressing TMPRSS2 (VTN cells). Critically we found that the ORF10KO virus is less pathogenic in a hamster model of SARS-CoV-2 infection and that the ORF10KO virus frequently reverts to WT during the course of infection.

## Materials and Methods

### Cell culture and virus propagation

Human A549 cells modified to constitutively express ACE2 and TMPRSS2 (A549-AT)(24) were a kind gift from Dr Suzannah Rihn, MRC-University of Glasgow Centre for Virus Research. An African green monkey kidney cell line (Vero E6) modified to constitutively express the human serine protease TMPRSS2 (Vero E6/TMPRSS2 (VTN), (25)) was obtained from NIBSC, UK. Both lines were maintained in DMEM supplemented with 100 U/ml penicillin and 100 µg/ml streptomycin and 10% foetal bovine serum. Recombinant virus stocks were grown in VTN cells with a starting multiplicity of infection (moi) of 0.01 and the infection allowed to proceed until extensive cell death was observed (usually 2-3 days post infection). The supernatant was then clarified by filtration, aliquoted and stored at -70 °C.

### Design and isolation of WT and ORF10KO recombinant SARS-CoV-2 viruses

Recombinant viruses were made using Transformation-associated Recombination in yeast (26) (TAR) as previously described (27). The recombinant wild type (WT) genomic sequence was constructed using the Wuhan-Hu-1 (GenBank accession: <u>NC_045512</u>) genome but modified to include the D614G mutation in the S protein. The ORF10KO recombinant virus was similarly made but overlapping PCR mutagenesis was used to change the initiating start codon of ORF10 from ATG to TAA and to change the internal methionine codon from ATG to TAG.

Briefly, SARS-CoV-2 replicon cDNA fragments were assembled into full-length cDNA clones by TAR assembly using the GeneArt™ High-Order Genetic Assembly System (Invitrogen™, ThermoFisher) according to the manufacturer’s instructions. Prospective clones were amplicon sequenced (NEBNext® ARTIC SARS-CoV-2 FS Library Prep Kit) prior to *in-vitro* generation of genomic RNA using T7 polymerase. T7 polymerase promoter driven *in-vitro* RNA transcription (IVT) from full length genome BAC/YAC clones and a construct containing a codon optimised SARS-CoV-2 N gene sequence, under control of a T7 promoter, were performed using a RiboMAX™ Large Scale RNA Production System-T7 (Promega) with the inclusion of m7G(5’)G RNA Cap Analog (NEB). Electroporation of the resulting RNA was carried out using a Neon® Transfection System (Invitrogen™, ThermoFisher) following the manufacturer’s protocol for adherent cell lines. Both rescued viruses were amplified in VTN cells and stocks titrated by TCID_50_ method as previously described (2).

### RNA isolation and amplification of the ORF10 coding region for Sanger Sequencing

Viral RNA from infected cell supernatants was extracted using a QIAamp Viral RNA Mini Kit (Qiagen) according to the manufacturer’s instructions. An 800 bp fragment covering the ORF10 region was amplified by one-step RT-PCR using the SuperScript™ IV One-Step RT-PCR System (Invitrogen™, ThermoFisher) and the following primers (OL11F: 5’-CTTTGCTGCTGCTTGACAGATTGAACCAGC-3’ and Wu30R: 5’-GGTGGCTCTTTCAAGTCCTC-3’). The PCR fragment was commercially sequenced (Eurofins) using the OL11F primer.

### RNA isolation and direct RNA sequencing

Approximately 10^7^ VTN cells were infected at a moi of 0.01 and incubated at 37 °C until substantial cytopathic effect (CPE) was observed (usually 2-3 days). Total RNA was extracted from the cells using Trizol as per manufacturer’s instructions but with 3 washes of the RNA pellet with 70% ethanol. Direct RNA sequencing was performed on a MinIT sequencing device (Oxford Nanopore Technologies, ONT) using the ONT direct RNA sequencing kit (SQK-RNA002) as per manufacturer’s instructions and the data was analysed as previously described (2). To increase the accuracy of mapping, WT virus and ORF10KO virus derived reads were aligned to bespoke FASTA files with a genome reflecting the expected sequence of each virus. Thus, the reads derived from WT infected cells were aligned to a Wuhan-Hu-1 genome containing the D614G point mutations (WT target genome). Reads derived from ORF10KO infected cells were aligned to a Wuhan-Hu-1 genome containing the D614G point mutations alongside the ATG >TAA and ATG>TAG mutations in the ORF10 region (ORF10KO target genome).

### RNA isolation and amplicon based sequencing

After sacrificing both groups of infected hamsters at 6 days post infection, the lungs were removed and total RNA was isolated by homogenising 0.1 g of lung tissue in a TissueLyser LT (Qiagen) then incubating with TRIzol™ (Invitrogen™, ThermoFisher) and extracting as per manufacturers instructions but with three washes of the RNA pellet with 70% ethanol. RNA samples were then used in the Midnight amplicon sequencing protocol from ONT as described by the manufacturer and the sequencing data was aligned to the SARS-CoV-2 genome using minimap2 (28). As with the direct RNA sequencing, we aligned the reads from the WT virus infected groups to the WT target genome. Similarly, we aligned the reads from the ORF10KO group to the ORF10KO target genome. Analysis of the data was done by an in-house script that examines each aligned sequence read in turn to determine the reported sequence for that read across three locations. Firstly, nucleotides 29556 – 29562 (AG AUG GG on the WT genome) which covers the initiating methionine for ORF10. Second, nucleotides 29602 – 29608 (U CUA CUC) which covers the region proposed to basepair with the initiating codon region in the RNA structure described (Figure 1b). Finally nucleotides 29615 – 29626 (AGA AUG AAU UCT) which cover the internal methionine codon and a surrounding region that may form a stem loop structure with this internal methionine codon. For each sequence read, the observed nucleotides across all three regions is recorded and in this way the script records how many times any particular pattern of nucleotides across the three regions is observed.

**Figure 1.**
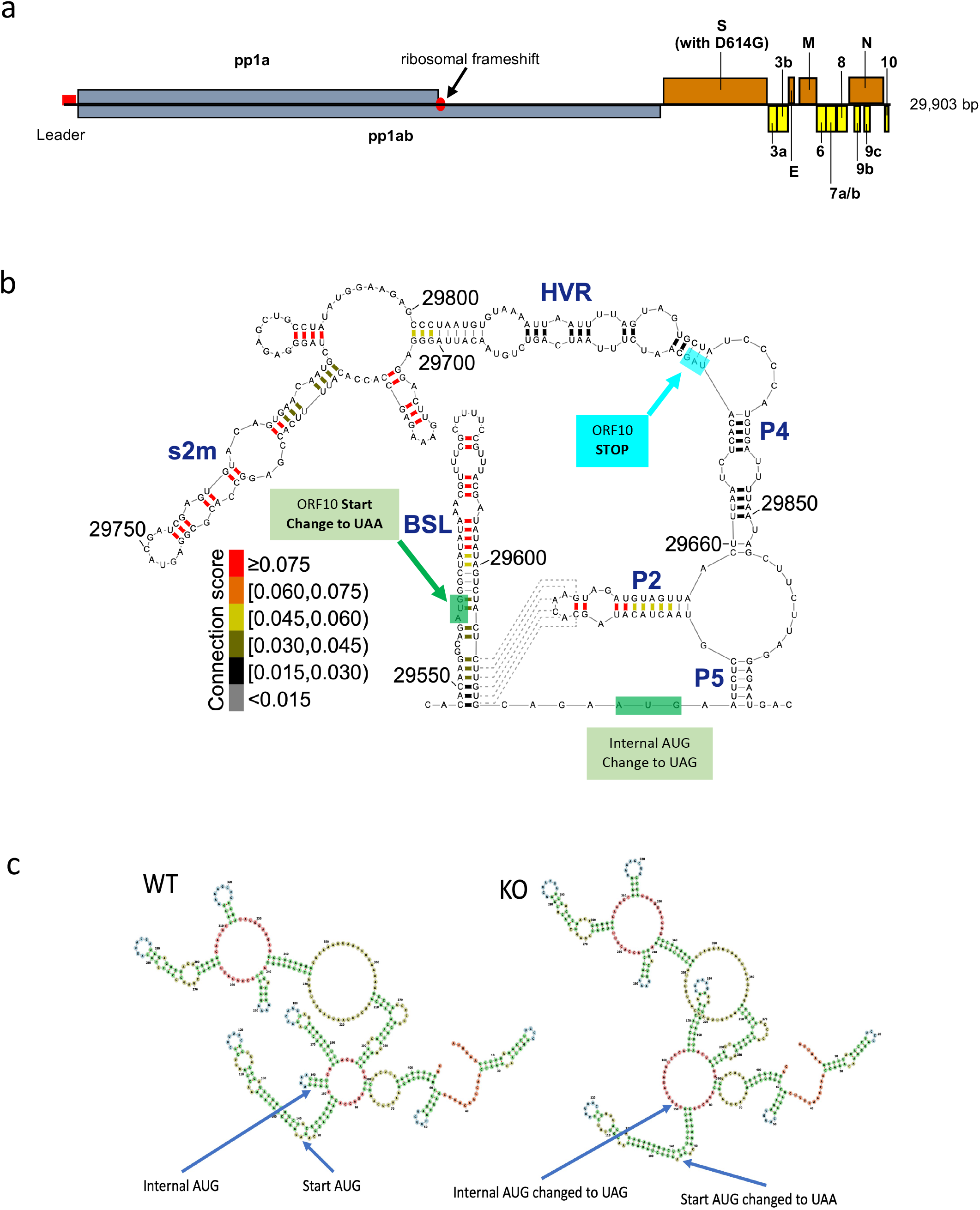
Overview of ORF10 on the SARS-CoV-2 genome, proposed RNA structures and properties of the ORF10KO virus. A simplified layout of SARS-CoV-2 genes is shown in part (a) with pp1a and pp1ab in grey, structural proteins in orange and non-structural accessory proteins in yellow. (b) Proposed structure of the 3’ UTR of SARS-CoV-2 taken from Cao et al.(15) and adapted to indicate clearly the locations of the point mutations introduced into the ORF10KO virus. Part (c) illustrates the predicted folding of the ORF10 region for WT and the ORF10KO virus using MXfold (32). The locations of the initiating and internal AUG codons are indicated.

In addition, for each of the 12 lung samples, we used RT-PCR to amplify just the ORF10 coding region as described earlier for sanger sequencing of cell culture derived virus. These PCR fragments were then barcoded (sequencing kit SQK-NBD114.24, ONT) and sequenced according to the manufacturer’s instructions using the MinIT device from ONT. The sequence data was aligned using minimap2 to appropriate target genomes - WT genome for samples from WT infected hamsters and ORF10KO genome for samples from ORF10KO infected hamsters. As before we analysed the alignments using an in-house script to catalogue and quantify sequences in the three regions described above. This approach proved more reliable in providing large numbers of sequence reads that unambiguously covered the ORF10 coding region.

### Passage of virus in cell culture for competition assay

Approximately 3×10^6^ cells were infected at an moi of 0.01 with a 50:50 mixture of the recombinant WT and ORF10KO viruses. A sample of the virus mixture (referred to as P0) was retained for Sanger sequencing. Cells were incubated for 48 hours before the supernatant was harvested and viral release was assayed by qRT-PCR as previously described (29). The harvested virus was then diluted as appropriate in order to re-infect another batch of 3×10^6^ cells and this process was repeated until the mixture of viruses had three passages in the cell lines required. After the final passage the viral RNA (referred to as P3) was extracted from the supernatant and the ORF10 region amplified by RT-PCR and sent for Sanger sequencing alongside the corresponding product from the P0 control sample.

### Proteomics analysis

Nine flasks of A549-AT cells each containing 10^7^ cells were mock infected, infected with ORF10KO or the WT control in triplicate. The infected cells were harvested at 16 hours post infection and processed for quantitative proteomics.

### TMT Labelling, High pH reversed-phase chromatography and Phospho-peptide enrichment

Aliquots of 100 µg of each sample were digested with trypsin (2.5 µg trypsin; 37 °C, overnight), labelled with Tandem Mass Tag (TMT) ten plex reagents according to the manufacturer’s protocol (Thermo Fisher Scientific, Loughborough, UK) and the labelled samples pooled.

For the Total proteome analysis, an aliquot of 50 µg of the pooled sample was desalted using a SepPak cartridge according to the manufacturer’s instructions (Waters, Milford, Massachusetts, USA). Eluate from the SepPak cartridge was evaporated to dryness and resuspended in buffer A (20 mM ammonium hydroxide, pH 10) prior to fractionation by high pH reversed-phase chromatography using an Ultimate 3000 liquid chromatography system (Thermo Fisher Scientific). In brief, the sample was loaded onto an XBridge BEH C18 Column (130Å, 3.5 µm, 2.1 mm X 150 mm, Waters, UK) in buffer A and peptides eluted with an increasing gradient of buffer B (20 mM ammonium hydroxide in acetonitrile, pH 10) from 0-95% over 60 minutes. The resulting fractions (15 in total) were evaporated to dryness and resuspended in 1% formic acid prior to analysis by nano-LC MSMS using an Orbitrap Fusion Tribrid mass spectrometer (Thermo Fisher Scientific).

For the Phospho proteome analysis, the remainder of the TMT-labelled pooled sample was desalted using a SepPak cartridge (Waters, Milford, Massachusetts, USA). Eluate from the SepPak cartridge was evaporated to dryness and subjected to TiO2-based phosphopeptide enrichment according to the manufacturer’s instructions (Pierce). The flow-through and washes from the TiO2-based enrichment were then subjected to FeNTA-based phosphopeptide enrichment according to the manufacturer’s instructions (Pierce). The enriched phosphopeptides were evaporated to dryness and then resuspended in 1% formic acid prior to analysis by nano-LC MSMS using an Orbitrap Fusion Tribrid mass spectrometer (Thermo Fisher Scientific).

### Nano-LC Mass Spectrometry

High pH RP fractions (Total proteome analysis) or the phospho-enriched fractions (Phospho-proteome analysis) were further fractionated using an Ultimate 3000 nano-LC system in line with an Orbitrap Fusion Tribrid mass spectrometer (Thermo Fisher Scientific). In brief, peptides in 1% (vol/vol) formic acid were injected onto an Acclaim PepMap C18 nano-trap column (Thermo Fisher Scientific). After washing with 0.5% (vol/vol) acetonitrile 0.1% (vol/vol) formic acid peptides were resolved on a 250 mm × 75 μm Acclaim PepMap C18 reverse phase analytical column (Thermo Fisher Scientific) over a 150 min organic gradient, using 7 gradient segments (1-6% solvent B over 1min., 6-15% B over 58min., 15-32%B over 58min., 32-40%B over 5min., 40-90%B over 1min., held at 90%B for 6min and then reduced to 1%B over 1min.) with a flow rate of 300 nl min^−1^. Solvent A was 0.1% formic acid and Solvent B was aqueous 80% acetonitrile in 0.1% formic acid. Peptides were ionized by nano-electrospray ionization at 2.0kV using a stainless-steel emitter with an internal diameter of 30 μm (Thermo Fisher Scientific) and a capillary temperature of 275°C.

All spectra were acquired using an Orbitrap Fusion Tribrid mass spectrometer controlled by Xcalibur 2.1 software (Thermo Fisher Scientific) and operated in data-dependent acquisition mode using an SPS-MS3 workflow. FTMS1 spectra were collected at a resolution of 120 000, with an automatic gain control (AGC) target of 200 000 and a max injection time of 50ms.

Precursors were filtered with an intensity threshold of 5000, according to charge state (to include charge states 2-7) and with monoisotopic peak determination set to peptide. Previously interrogated precursors were excluded using a dynamic window (60s +/-10ppm). The MS2 precursors were isolated with a quadrupole isolation window of 1.2m/z. ITMS2 spectra were collected with an AGC target of 10 000, max injection time of 70ms and CID collision energy of 35%.

For FTMS3 analysis, the Orbitrap was operated at 50 000 resolution with an AGC target of 50 000 and a max injection time of 105ms. Precursors were fragmented by high energy collision dissociation (HCD) at a normalised collision energy of 60% to ensure maximal TMT reporter ion yield. Synchronous Precursor Selection (SPS) was enabled to include up to 10 MS2 fragment ions in the FTMS3 scan.

### Proteomics Data Analysis

The raw data files were processed and quantified using Proteome Discoverer software v2.4 (Thermo Fisher Scientific) and searched against the UniProt Human database (downloaded January 2023: 81579 entries) and the SARS-CoV-2 protein list using the SEQUEST HT algorithm. Peptide precursor mass tolerance was set at 10ppm, and MS/MS tolerance was set at 0.6Da. Search criteria included oxidation of methionine (+15.995Da), acetylation of the protein N-terminus (+42.011Da) and methionine loss plus acetylation of the protein N-terminus (-89.03Da) as variable modifications and carbamidomethylation of cysteine (+57.0214) and the addition of the TMT mass tag (+229.163) to peptide N-termini and lysine as fixed modifications. For the Phospho-proteome analysis, phosphorylation of serine, threonine and tyrosine (+79.966) was also included as a variable modification. Searches were performed with full tryptic digestion and a maximum of 2 missed cleavages were allowed. The reverse database search option was enabled and all data was filtered to satisfy false discovery rate (FDR) of 5%.

### Pathogenesis of viruses in the Syrian hamster model

Animal work was approved by the local University of Liverpool Animal Welfare and Ethical Review Body and performed under UK Home Office Project Licence PP4715265. Male Syrian Golden hamsters (8-10 weeks old) were purchased from Janvier Labs (France). Animals were maintained under SPF barrier conditions in individually ventilated cages. Animals were randomly assigned into two cohorts of 6 animals. For SARS-CoV-2 infection, hamsters were anaesthetised lightly with isoflurane and inoculated intra-nasally with 100 µl containing 10^4^ PFU SARS-CoV-2 (either ORF10KO or WT rescue) in PBS. Weights were recorded daily and oral swabs were obtained under isoflurane anaesthetic at 3 and 5 dpi for quantification of viral load by qRT-PCR. Animals were sacrificed at 6 days post-infection by an overdose of pentabarbitone. Tissues were removed immediately for downstream processing.

For quantification of viral load, RNA was extracted from swab and tissue samples as described above and qRT-PCR using primers directed against the N gene were performed as described previously (30). Quantification was determined using a standard curve and values normalised relative to 18S RNA.

From all animals the left lung was fixed in 10% buffered formalin for 48 h and then stored in 70% ethanol until further processing. The lung was halved by a longitudinal section and routinely paraffin wax embedded. Consecutive sections (3-5 µm) were prepared and stained with hematoxylin eosin (HE) for histological examination or subjected to immunohistological staining. Immunohistology was performed to detect SARS-CoV-2 antigen, using the horseradish peroxidase (HRP) method and the following primary antibody: rabbit anti-SARS-CoV nucleocapsid protein (Rockland, 200-402-A50) as previously described (31). For quantification of SARS-CoV-2 antigen expression and the proportion of non-aerated parenchyma vs aerated parenchyma, a morphometric analysis was undertaken on the sections stained for SARS-CoV-2 NP and by HE, respectively. The stained slides were scanned using NanoZoomer 2.0-HT (Hamamatsu, Hamamatsu City, Japan), and the lung sections from each animal were quantitatively analysed using the Visiopharm 2022.01.3.12053 software (Visiopharm, Hoersholm, Denmark). In HE-stained section, the morphometric analysis served to quantify the area of non-aerated parenchyma and aerated parenchyma in relation to the total area (= area occupied by lung parenchyma on two sections prepared from the left lung lobes) in the sections, as previously described (31). For each section stained for SARS-CoV-2 NP, the lung was manually outlined in Visiopharm, and annotated as a Region Of Interest (ROI), manually excluding artifactually altered areas. The manual tissue selection was further refined with an Analysis Protocol Package (APP) based on a Decision Forest classifier, with the pixels from the ROI being ultimately classified as either “Tissue” or “Background”. This new “Tissue” ROI, regrouping the lung samples analysed for each animal, was further quantified by executing two APPs successively. The first APP was based on a Threshold classifier and served to detect and outline areas with immunostaining. The second APP then measured both the surface of the immunostained area (µm2) and the surface of the “Tissue” ROI (µm^2^). The percentage of immunostained area (%), expressed as the ratio between the immunostained area and the total area, was obtained for each animal in Excel (Microsoft Office 2019; Microsoft, Redmond, Washington, United States), according to the following formula: ([positive area (µm2)]/ [total area (µm2)]) x 100.

### Data and code availability

Raw sequencing data (fastq files), genome sequences and in house analysis script is available from Zenodo (https://doi.org/10.5281/zenodo.8139581).

## Results

### Viable rescue of ORF10KO virus and WT control

Figure 1a illustrates the location of ORF10 at the 3’ end of the viral genome and illustrates the change of the sequence of the original Wuhan virus spike protein to one with the D614G mutation. Figures 1b illustrates the predicted folding of the viral RNA present in this region (15) and shows the mutations made to generate the ORF10KO virus with the initial start codon changed to UAA (all three nucleotides changed) as well as the second internal AUG changed to UAG (two nucleotide changes). Thus, for such a virus to revert to wild type there would be five nucleotide changes needed, and at least four changes to regain an open reading frame (three to gain a start codon and at least one more to remove the introduced internal stop codon). Although the predicted RNA structure in Figure 1b implies that the internal AUG is not in a structured part of the viral genome, RNA structure prediction approaches do vary in their conclusions. Figure 1c shows structure predictions made by MXfold (32) and illustrates the potential for the introduced changes to affect viral RNA folding in this region.

Rescue of the ORF10KO virus and the WT counterpart was successful in both cases. We also attempted to rescue two reporter viruses, one with ORF10 replaced with the mScarlet coding sequence and a second virus with ORF10 replaced with the mNeonGreen coding sequence. We were unable to rescue either virus despite three attempts at each virus rescue.

To analyse the transcriptome of the two rescued viruses, VTN and A549-AT cells were separately infected with the two viruses and the infected cell mRNA was harvested prior to sequencing by direct RNA sequencing (dRNAseq) on Oxford Nanopore. The mRNA sequence reads were mapped to the viral genome using minimap2(28). Figure 2a illustrates the read depth for mRNA transcripts across the viral genome for both the WT and ORF10KO viruses and Figure 2b shows both direct RNA sequence data and Illumina amplicon sequence data (derived from the VTN cells) across the region where the stop codons were introduced.

**Figure 2.**
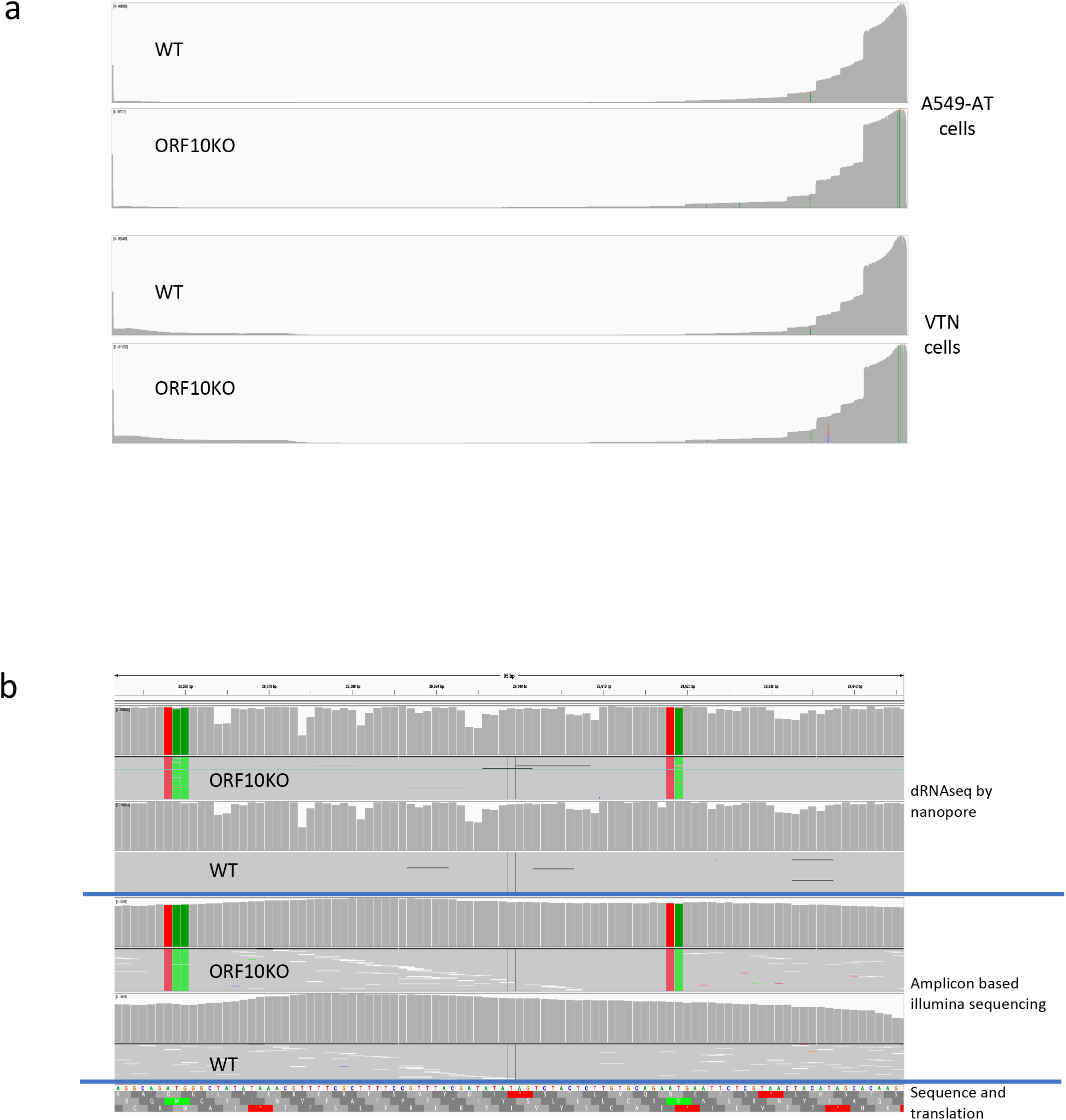
Transcriptome and sequences of WT and ORF10KO viruses. Part (a) shows total viral mRNA sequenced by direct RNA sequencing of mRNA extracted from A549-AT cells and VTN cells infected with ORF10KO or WT virus and mapped to the SARS-CoV-2 genome, image taken from IGV viewer. Part (b) shows an IGV viewer snapshot of sequencing data focusing on the ORF10 region. The top two sections are from direct RNA sequencing of mRNA extracted from VTN cells. The bottom two sections show the results of Illumina based amplicon sequencing of RNA extracted from the supernatants of the two rescued viruses. The coloured lines indicate changes from the WT sequence and are as expected.

### Growth rates of the WT and ORF10KO viruses in different cell lines

The growth kinetics of the two viruses were compared in two cell lines with distinct properties that are commonly used to grow SARS-CoV-2 (Figure 3). The A549-AT cell line is a human lung carcinoma cell line engineered to over express ACE2 and TMPRSS2 and has a complete innate immune gene repertoire. In this cell line we noted an apparent small growth advantage for the ORF10KO virus but this slight advantage was not statistically significant over three repeats. The VTN cell line is derived from VeroE6, a non-human primate cell line with a defect in the innate immune response due to loss of the type 1 interferon gene cluster(33). In VTN cells the two virus’s growth curves are very closely matched and appear identical by visual inspection.

**Figure 3.**
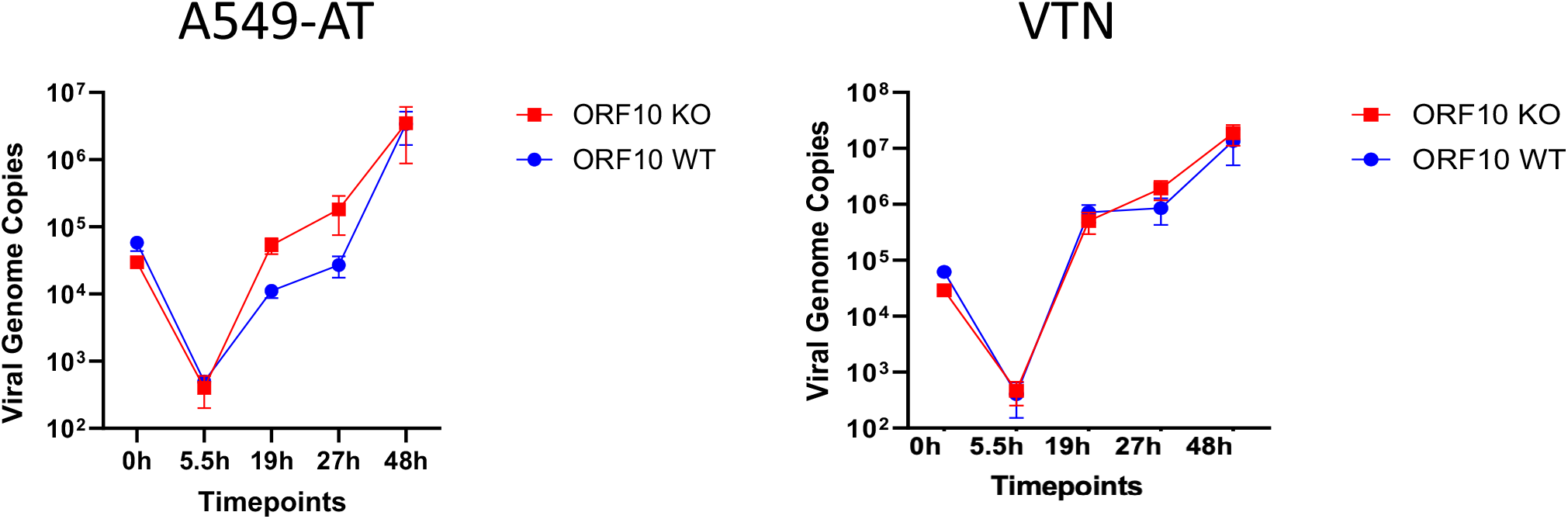
Viral growth curves in VTN and A549-AT cells. Both VTN and A549-AT cell lines were used to determine the relative growth of the WT (blue) and ORF10KO (red) viruses. The amounts of viral RNA in the culture supernatants were assayed by qRT-PCR at the time points indicated, in triplicate.

In order to explore further the possibility that the two viruses have different growth advantages in different cell lines we used a competition experiment where the two viruses are passaged together and the resulting virus mix is sequenced to determine if either virus has a growth advantage. Figure 4 shows the result of independent growth competition assays in A549-AT and VTN lines. Here we see that in A549-AT cells, compared to the sequence data from the mixed inoculum, the ORF10KO virus either totally dominates (replicate 1) or out competes the WT virus (replicates 2-3). In contrast, in VTN cells the WT virus clearly dominates in each competition experiment. This is despite the observation from the individual virus growth curves that the two viruses have apparently indistinguishable growth properties in this cell line.

**Figure 4.**
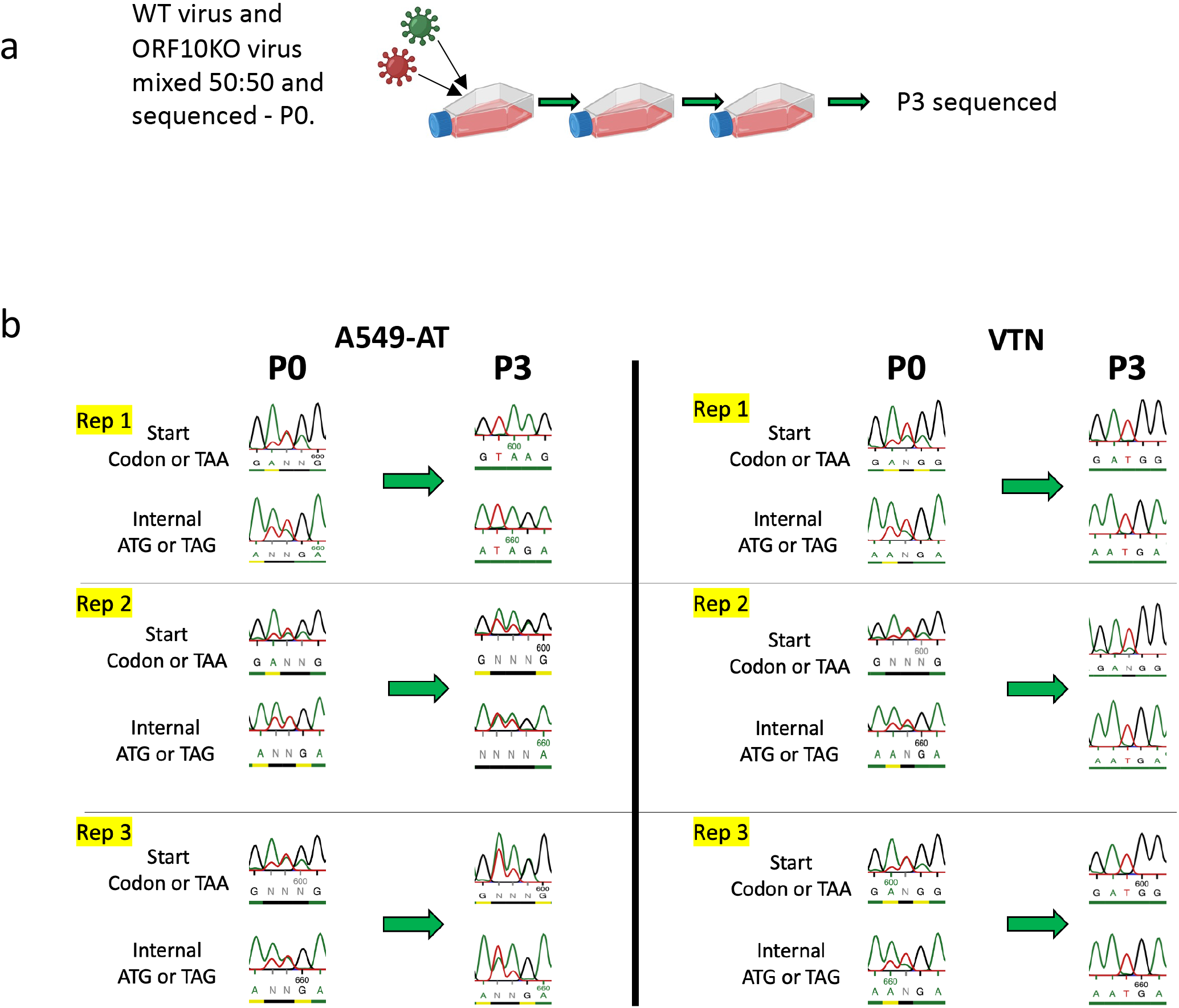
Co-infection competition in two cell lines. Part (a) illustrates the experimental set up. A 50:50 mix of the WT and ORF10KO viruses was used to infect either VTN or A549-AT cells at an moi of 0.01. After 48 hours a sample of the supernatant was removed, assayed by qRT-PCR and diluted as appropriate to infect another flask of cells again at an moi of 0.01. Samples from both the input and passage 3 supernatants were taken, the RNA extracted and the ORF10 region was reverse transcribed and PCR amplified prior to Sanger sequencing to determine if one or other virus was able to dominate after three passages. The relevant regions of the Sanger sequencing data is shown (b) for each of the three biological replicates undertaken in each cell line.

### Virus growth competition in the presence of Ruxolitinib, a JAK/STAT pathway inhibitor

A key difference between VeroE6 and A549 based cells is that VeroE6 cells have an ablated IFN type I gene cluster (34). In contrast this is not the case for A549 cells and so we wanted to test the possibility that innate signalling via the JAK/STAT pathway may affect the outcomes of the competition experiment. The competition experiment was therefore repeated in A549-AT cells in the presence of ruxolitinib, which inhibits JAK1/2 function(35), but this did not affect the outcome and the ORF10KO virus still dominated after 3 passages (Supplementary Figure 1). Western blot analysis of A549-AT cells with or without ruxolitnib treatment using antisera against STAT1 and phosphorylated STAT1 showed that the treatment successfully inhibited STAT phosphorylation (data not shown).

### Proteomics analysis of A549-AT cells infected with WT or ORF10KO

To study the host cell response to infection with the WT and ORF10KO viruses, A549-AT cells were infected at an moi of 3 with either the WT or ORF10KO virus or mock infected. At 16 hours post-infection lysates were prepared from the infected cells and analysed by TMT labelling in combination with LC-MS/MS. Figure 5 shows a volcano plot of protein abundance changes in WT vs ORF10 infected cells illustrating that both viruses affect the host cell proteome similarly. Indeed, we detected very few proteins whose abundance was differentially affected (e.g. a larger than 2-fold change) by the two viruses relative to each other (Supplementary Table 1).

**Figure 5.**
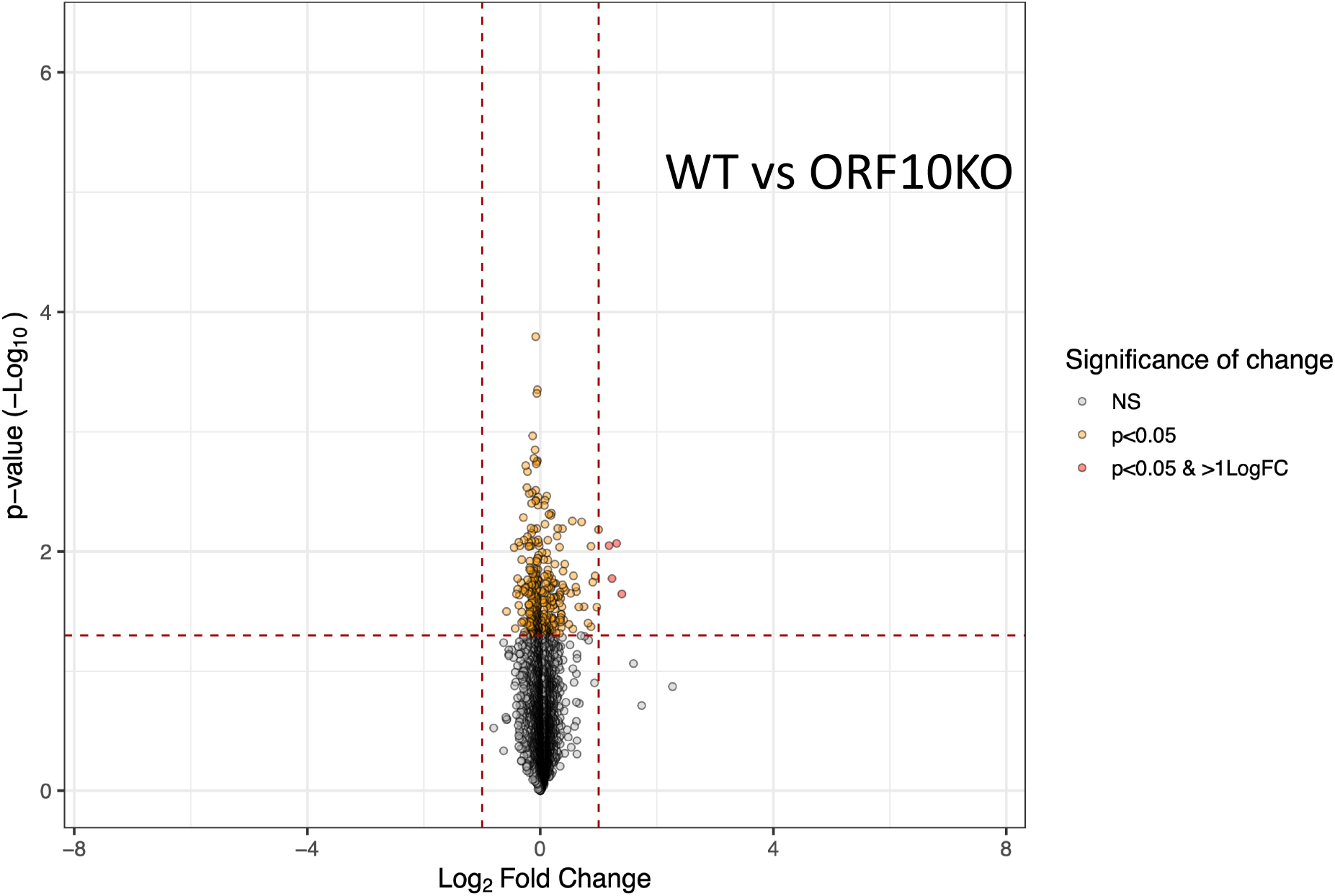
Proteomics analysis of ORf10KO vs WT in A549-A2 cells. Volcano plot illustrating the changes in abundance of cellular proteins alongside the p-values in cells infected with either WT or ORF10KO virus.

### Infection of Syrian golden hamsters with the WT and ORF10KO viruses reveals distinct pathogenicity and growth characteristics in the upper and lower respiratory tracts

To determine the relative fitness and pathogenicity of the ORF10KO and WT rescue virus, two groups of six hamsters were infected intranasally with 10^4^ pfu of either the WT virus (WT group) or the ORF10KO virus (ORF10KO group). Weight loss is a clinical measure of severity of disease in the hamster model. Over 6 days, hamsters infected with the WT virus lost approx. 15% of their body weight which is typical for the WT strain of SARS-CoV-2 (Fig. 6a). However, animals infected with the KO virus lost significantly less weight (6%; P < 0.01, 2-way ANOVA with Bonferroni post-test). Assessment of the viral RNA load in throat swabs taken at 3 and 5 days post infection revealed that there was no significant difference in the viral load between WT and ORF10KO at both time points (Fig. 6b). All animals were sacrificed at day 6 post-infection and viral load in nasal mucosa and lung samples was quantified by qRT-PCR. The results (Fig 6c) showed there was no significant difference between the groups in nasal tissue. However, there was an approximately 1 log reduction in viral load in the lungs of ORF10KO virus-infected animals compared to WT virus infected animals (P < 0.01, Mann-Whitney *U* test, two-tailed). The histological examination of the lungs showed that in both groups there were changes consistent with SARS-CoV-2 infection in hamsters at 6 dpi. That is, the parenchyma exhibited coalescing consolidated areas with activated and hyperplastic type II cells, some desquamed alveolar epithelial cells and an inflammatory infiltrate composed of macrophages, lymphocytes, and fewer neutrophils. Morphometric analysis of the lungs to objectively quantify the area occupied by aerated and consolidated, i.e., non-aerated parenchyma confirmed that the extent of parenchymal consolidation was significantly lower in ORF10KO as compared with WT animals (Fig. 6e; P < 0.01, Mann-Whitney *U* test, two-tailed), consistent with a less intense inflammatory response. Immunohistological examination (detection of SARS-CoV-2 NP) of lungs showed viral antigen expression in alveolar epithelial cells and some macrophages within focal areas of consolidation in both groups (Fig 6d). Morphometric analysis to quantify the extent of antigen expression showed significantly less staining in animals infected with the ORF10 virus (*P* < 0.01, Mann-Whitney *U* test, two-tailed). Thus, the ORF10 KO virus is less pathogenic, with lower viral loads in the lungs at 6 dpi.

**Figure 6.**
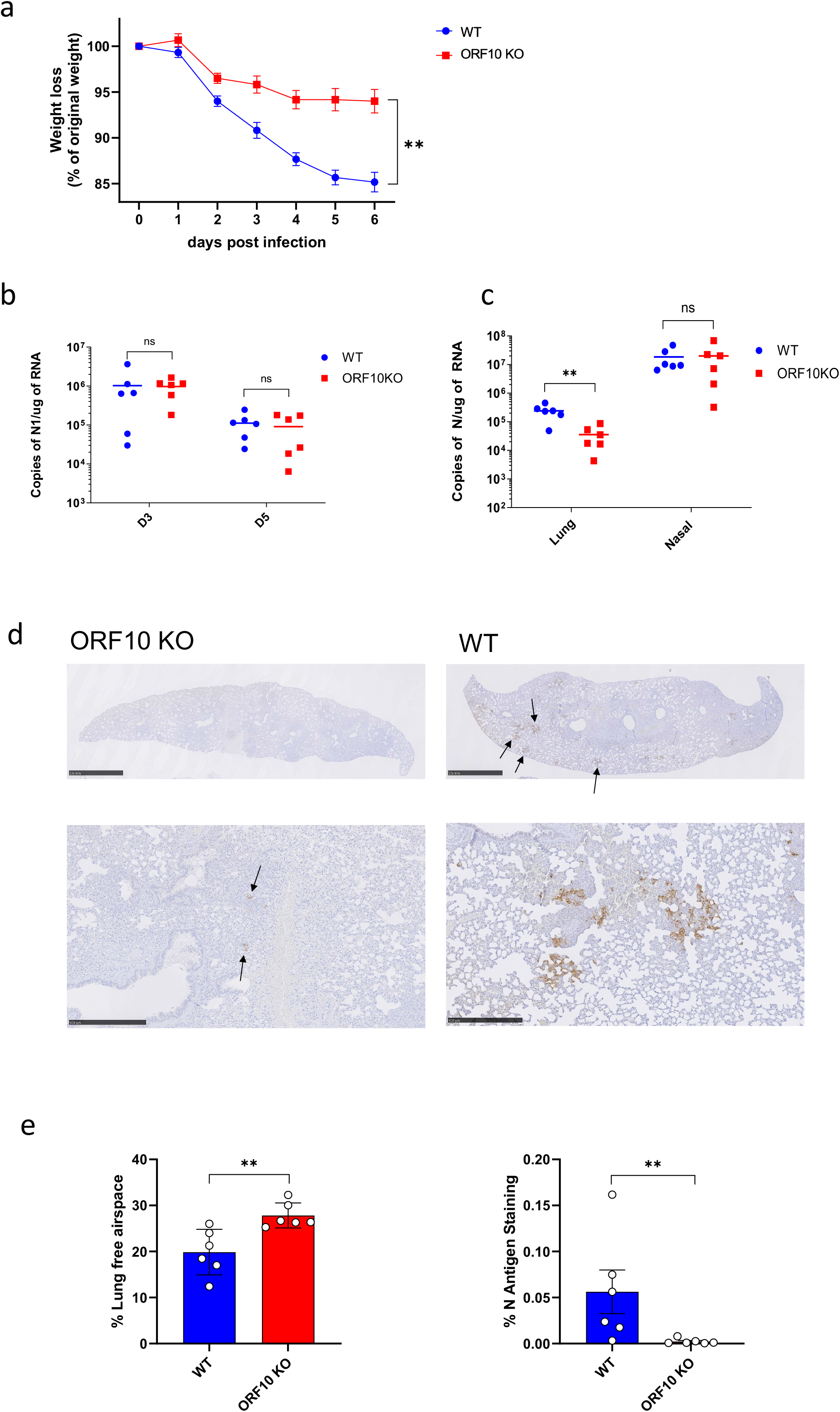
ORF10KO virus is attenuated in a hamster model of infection. Two groups of six male Golden Syrian hamsters (8-10 weeks old) were infected with 10^4^ PFU of either the WT or ORF10KO virus. (a) Hamsters were monitored for weight loss at indicated time-points. Points show the mean ± SEM. Curves were analysed by two-way ANOVA with Bonferroni post-tests. SARS-CoV-2 viral load in throat swabs at days 3 and 5 (b) and in nasal and lung tissues (c) was determined using qRT-PCR for the N gene. Points represent individual animals with a line at the mean. Side-by-side comparisons were made using Mann-Whitney U test. ** represents *P* < 0.01 and ns non-significant. (d) Representative images of lung sections stained by immunohistology for SARS-CoV N antigen (brown), using a rabbit polyclonal antibody, with haematoxylin counterstain. Upper panels show a whole lung section (scale bars represent 2.5 mm) and lower panels are a higher magnification image taken from the same section (scale bars represent 500 µm). The extent of viral antigen expression in the ORF10KO-infected hamster lung is much lower compared to a WT-infected hamster lung. Arrows point to areas of antigen staining. (e) Morphometric analysis of scanned HE-stained sections was performed using the software programme Visiopharm to quantify the area of non-aerated parenchyma and aerated parenchyma in relation to the total area. Results are expressed as the mean free airspace in lung sections (left panel). A similar morphometric analysis was performed to quantify the area stained for SARS-CoV-2 N antigen (right panel). In ORF10KO-infected hamster lungs, the area of aerated parenchyma is significantly higher, and the area of viral antigen expression significantly lower, indicating less extensive lung infection and a less intense inflammatory response. Points represent individual animals with bar at the mean. Pairwise comparisons were made between groups using a Mann-Whitney U test. ** represents *P* < 0.01.).

### Amplicon sequencing of virus genome in lung tissue at day 6 reveals frequent reversion to WT in ORF10KO infected hamsters

RNA was extracted from the lungs of the 12 infected hamsters and subjected to nanopore based amplicon sequencing. The sequence data was mapped to the viral genome (for WT infected samples) or to an ORF10KO modified viral genome (for the ORF10KO infected samples) using minimap2 (28). The resulting aligned sequences were analysed using an in-house script that focussed on the initiating and internal AUG codons (changed to UAA and UAG respectively in the ORF10KO virus) as well as the region proposed to be complimentary to the initiating AUG. This script records and counts how often a sequence is observed at each codon and whether sequence reads map only to one of the three regions analysed or to two or three. There were a number of individual sequences within the dataset where the individual mapped sequence did not cover both the start and internal AUG. As a result we have reported the total number of times we observed either a start or a stop codon at each location as well as the number of times we observed a full ORF10KO or WT sequence on a continuous sequence. Tables 1 and 2 indicate how often the expected codons for WT vs ORF10KO were observed at the initiating or internal AUG within ORF10.

**Table 1.** Quantitative summary of observed mutations within the sequence data from lungs of hamsters infected with WT virus. Summary of observed sequences at the initiating AUG codon and the internal AUG codon for the WT virus infected hamster lungs. For each animal we counted total number of sequence reads with some sequence data at the two locations. In addition we noted how many times we observed either the exact AUG or UAA codon at the initiating codon and the exact AUG or UAG at the internal codon. We also noted how frequently there was a deletion of the G in the second observed codon. Finally, we also recorded how frequently we observed that both codons were either WT or ORF10KO matches on the same sequence read (i.e. indicating fully WT or fully ORF10KO).

|  |  |  |  |  |  | Entries<br>with<br>deletion of<br>G from<br>internal<br>AUG | entries<br>which<br>match<br>WT | entries<br>which<br>match<br>ORF10KO | Ratio<br>AUG/AUG<br>to<br>STOP/STOP |
| --- | --- | --- | --- | --- | --- | --- | --- | --- | --- |
| Name | entries with<br>seq data | entries with<br>exact initial<br>START | entries with<br>exact initial<br>STOP | entries<br>with exact<br>*internal*<br>AUG | entries<br>with exact<br>*internal*<br>STOP |  |  |  |  |
| WT1 | 3281 | 2537 | 0 | 1986 | 0 | 68 | 1550 | 0 | 0 |
| WT2 | 13281 | 10302 | 1 | 8229 | 0 | 240 | 6464 | 0 | 0 |
| WT3 | 9712 | 7454 | 0 | 5921 | 0 | 164 | 4558 | 0 | 0 |
| WT4 | 2997 | 2198 | 1 | 1731 | 2 | 46 | 1266 | 1 | 0.00078 |
| WT5 | 3096 | 2317 | 1 | 1920 | 1 | 67 | 1449 | 1 | 0.00069 |
| WT6 | 3010 | 2388 | 0 | 1823 | 0 | 56 | 1451 | 0 | 0 |

**Table 2.**
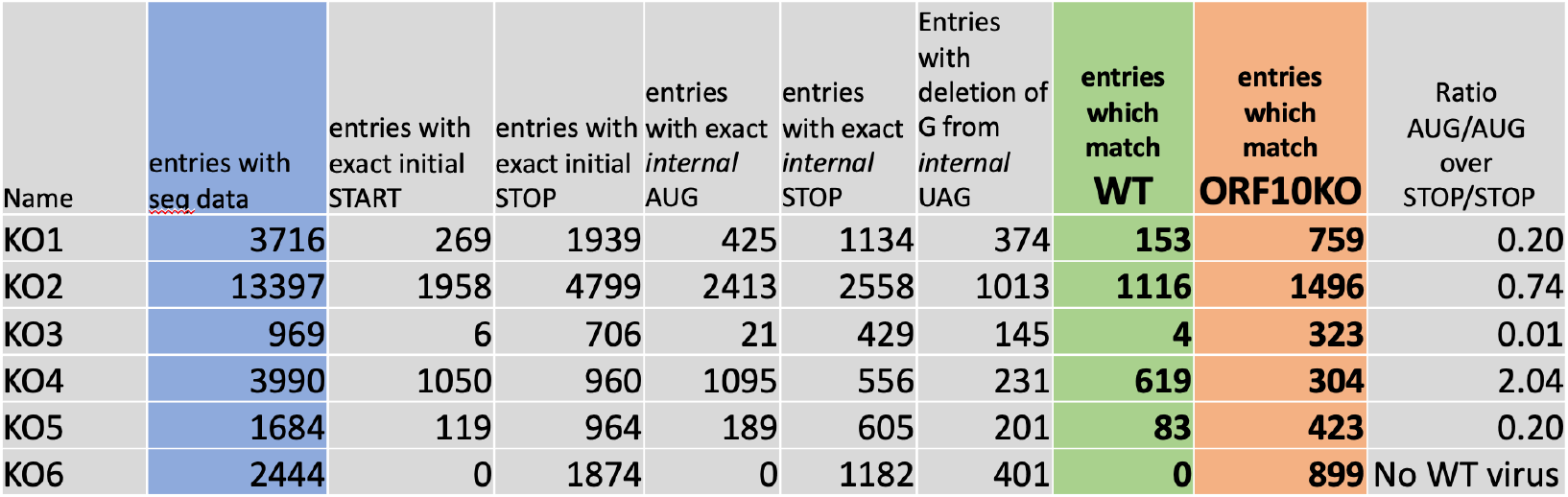
Quantitative summary of observed mutations within the sequence data from lungs of hamsters infected with ORF10KO virus. Summary of observed sequences at the initiating AUG codon and the internal AUG codon for the ORF10KO virus infected hamster lungs. For each animal we counted the total number of sequence reads with some sequence data at the two locations. In addition, we noted how many times we observed either the exact AUG or UAA codon at the initiating codon and the exact AUG or UAG at the internal codon. We also noted how frequently there was a deletion of the G in the second observed codon. Finally we also recorded how frequently we observed that both codons were either WT or ORF10KO matches on the same sequence read (i.e. indicating fully WT or fully ORF10KO).

Within the hamsters infected with the WT virus we observed that the sequences were relatively stable and that whilst it was possible to detect ORF10KO virus sequence in two of the hamster samples, it was rare (Table 1). The most common sequence change observed was a deletion of the G in the internal AUG (Supplementary Table 2). By comparison, in all but one of the hamsters infected with ORF10KO we were able to readily detect reversion to WT by at least one of the mutated codons and we were also able to show full reversion to WT (i.e. both codons reverted to AUG on the same sequenced molecule, Table 2). This reversion to WT ranged from rare (hamster KO3) to being more prevalent than the ORF10KO virus (hamster KO4, Table 2). As with the WT infected hamsters we also saw a frequent deletion of the internal G in the internal AUG (changed to UAG in the ORF10KO virus) which was sometimes the most frequent change (Table 2 and Supplementary Table 2). Visual inspection of the mapped reads across this region using IGV viewer correlated with these observations in both groups of animals (Figure 7a,b).

**Figure 7.**
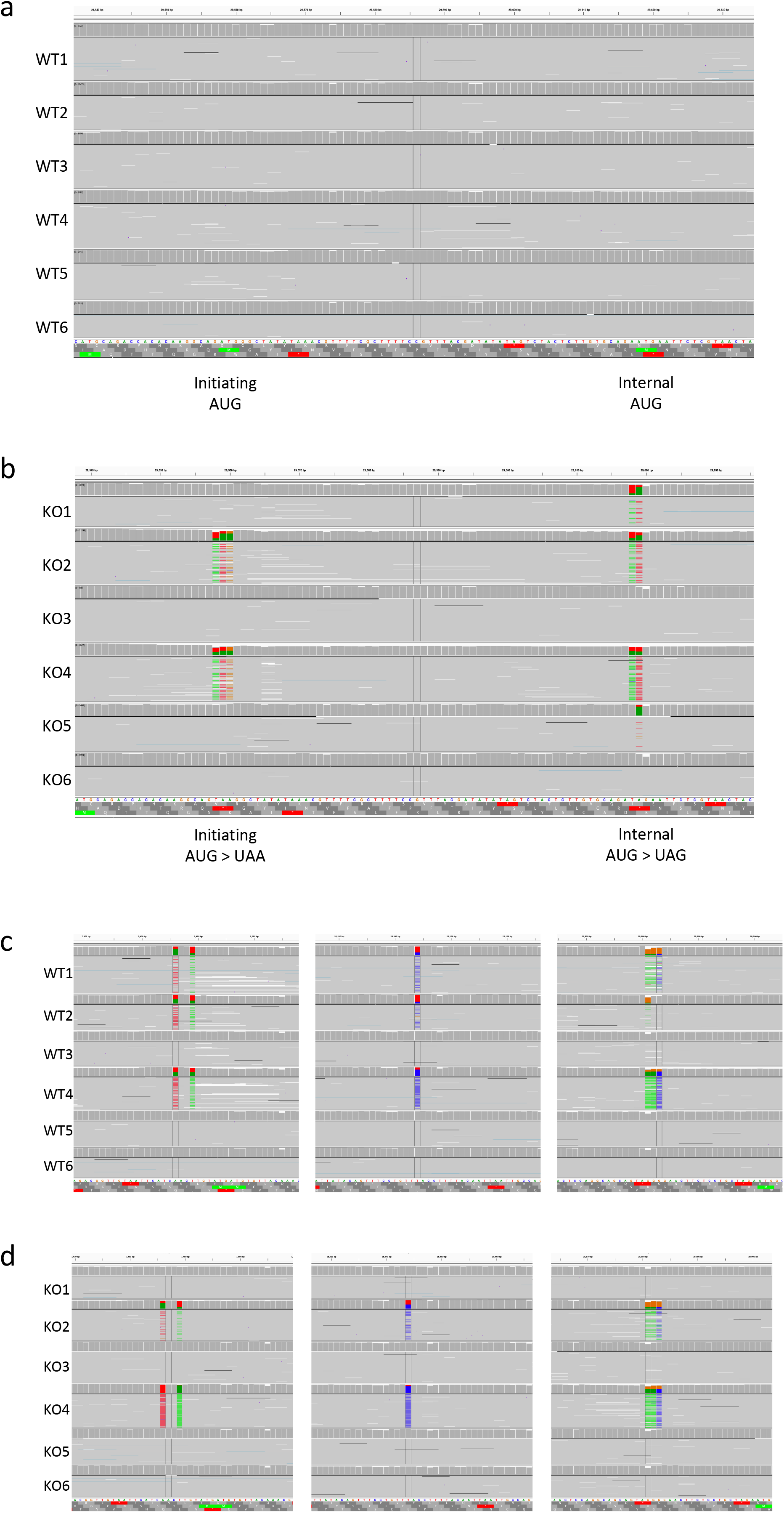
Alignments of amplicon based sequence data from viral genomes in lung tissues showing ORF10 region. RNA samples taken from the lungs of all the hamsters were amplicon sequenced and aligned to the relevant SARS-CoV-2 genome. Parts (a) and (b) show the alignments in the ORF10 region to illustrate that for the sequences derived from all six hamsters in the WT infection group (a) there is little variation from the expected WT sequence. However, examination of the same region on the ORF10KO virus genome (b) reveals that there is clear evidence of sequence deviation from the expected ORF10KO sequence (where both AUGs are altered) in some of the hamsters. Genomic sequence and all frames translation is shown at the bottom of both alignment figures, screen shot taken from IGV. Parts (c) and (d) illustrate examples of similar point mutations occurring elsewhere on the viral genome in samples taken from both groups of hamsters.

We noted that the whole genome amplicon sequencing approach was not always able to provide confident linkage between mutations at different sites along the ORF10 region. To provide an alternative independent analysis of the region we used our ORF10 region specific primers to RT-PCR amplify the region from each lung sample and subsequently sequenced the amplified DNA products using the ONT device. This analysis correlated with the whole genome amplicon data outcomes except that in this analysis we were able to detect fully reverted WT sequences in all six hamsters infected with ORF10KO virus (Supplementary Table 3).

### Analysis of possible second site mutations in ORF10KO infected hamsters

Analysis of the rest of the viral genome revealed several locations where additional mutations in the ORF10KO viral genome apparently correspond with increased reversion to WT in samples KO2 and KO4 (Figure 7d). These three locations are within pp1ab (nt 7485 A>U and nt 7489 U>A, both synonymous), ORF8 (nt 28144 U>C, synonymous) and N (nt 28,880 GGG > AAC changing R202K and G203R). There are other point mutations within the amplicon data but only these three appear to correlate with the reversion to WT in terms of relative abundance. Due to the nature of amplicon sequencing we cannot establish linkage with the reversion. Notably, analysis of the samples from hamsters infected with WT virus we found similar mutations (Figure 7a) indicating that these SNPs can occur independently of the reversion to WT seen in the hamsters infected with ORF10KO virus.

## Discussion

We made an ORF10KO virus and have shown that compared to WT this virus has distinct properties in cell culture and in the widely used hamster model of SARS-CoV-2 infection. Most strikingly we have shown that the ORF10KO virus is markedly less pathogenic in hamsters. Our data suggests that replication of the ORF10KO virus is restricted in the lower respiratory tract and that there is frequent reversion to WT during the course of a single infection. Allied to this we now have a reproducible cell culture based assay that allows us to select for the WT and ORF10KO viruses, showing that the two viruses have distinct growth properties in different cell lines. Taken together our data supports, but does not prove, the hypothesis that the ORF10 protein is expressed and functional *in vivo* and *in vitro*. Moreover, even if ORF10 protein is not required or functionally relevant, this data underlines the significance of the 3’ UTR in viral pathogenesis.

We reasoned that looking for reversion mutations in the ORF10KO virus genome *in vivo* would be highly informative as this allows for an unbiased selection of the most fit revertant in an *in vivo* environment. That we observe frequent reversion to WT ORF10 sequence does support the contention that ORF10 protein is functionally expressed and contributes to replication in the lower respiratory tract which in turn enhances the pathogenicity of SARS-CoV-2 in the hamster model. The recent suggestion by Khan *et al*. that ribosome binding within the ORF10 region leads to RNA folding changes within the 3’UTR (23) does partially align with our observations. However, there are some significant differences since our data suggests both AUG’s are selected for whereas Khan *et al*. proposed that the internal AUG is more important. Moreover, if ribosome binding is the critical factor, then single point mutations at either or both of the locations altered within the ORF10KO virus would produce an AUG at either location, albeit out of frame with the rest of putative ORF10.

Thus, a change in the ORF10KO virus from UAA GGC to U<u>AU G</u>GC at the start of ORF10 would restore a ribosome binding site. Similarly, changing AGA UAG to AG<u>A UG</u>G within the second mutated site would also provide a ribosome binding site. We did not observe either mutations in our sequencing data or indeed any other mutations that could affect RNA stem structure formation whilst retaining the ORF10KO sequence changes we introduced (Tables 1 and 2 and supplementary tables ST1 and ST2). There are a range of possible point mutations within and around the changes we made for the ORF10KO virus which would potentially restore stem-loop structures and/or ribosome binding sites. That full reversion is readily seen argues that either ORF10 protein expression is required and the protein product is functional or that the RNA sequence and structure formed is highly specific and does not tolerate any changes at these two locations. Notably, the second STOP codon we introduced into the ORF10 sequence reverted back to AUG and not to any other codon which would have opened the reading frame. This might suggest that if ORF10 protein is expressed and functional then this internal methionine is important to function and/or that there is a functional aspect of RNA folding here that does not tolerate change. What is also notable is that these experiments underline the potential for coronaviruses to acquire a number of point mutations within one infected individual if there is strong selective pressure. This rapid acquisition of multiple point mutations may be relevant to understanding the pace at which beneficial combinations of SNPs can arise and be selected for in the early stages of a zoonotic spill over.

We also observed a frequent single nucleotide deletion of G from the internal methionine codon within the ORF10 sequence, both in the samples sequenced from the WT virus infected hamster lungs and the ORF10KO counterparts. Modelling suggests this would have no effect on the predicted RNA structures for either virus and its significance is uncertain, this mutation is not present in the CoV-GLUE database (36) suggesting that such mutants cannot sustain person to person spread. This deletion would remove the internal methionine from a WT virus as well as change the reading frame of the putative ORF10 in a WT setting.

Other teams have deleted SARS-CoV-2 accessory proteins from the viral genome and shown reduced pathogenesis including, for example, ORF6 (37). Indeed, the differential replication we observe in the upper vs the lower respiratory tract has also been observed for SARS-CoV-2 viruses with deletions in the spike protein furin cleavage site (38, 39). One possibility is that this ORF10KO mutation could be combined with other deletions to generate a robust live attenuated vaccine strain either for SARS-CoV-2 or future coronaviruses with similar ORF10 like regions.

Several proteins have been proposed to be changed in abundance by ORF10 protein expression (12), but our proteomics data does not support a role for ORF10 protein in widespread changes in host cell protein abundance. Indeed, we find little evidence for differential protein abundance comparing WT and ORF10KO infected cells suggesting that the differences between the WT and ORF10KO viruses are mediated by more nuanced mechanisms.

That the ORF10KO virus has distinct growth characteristics in VeroE6 derived cells compared to A549 derived cells provides a useful platform with which to further study the properties of the ORF10KO virus. This observation may also hint at an explanation for the different growth characteristics in the hamster lung where ORF10KO virus growth appears to be restricted compared to nasal swabs where ORF10KO virus propagates as well as WT. We hypothesise that there could be one or more specific cell types in the lung within which the presence of a fully WT ORF10 region is highly advantageous.

Further work is underway to examine fully the role of the ORF10 region and to try to determine if the cell culture growth characteristics and *in vivo* pathogenesis is indeed a result of ORF10 protein expression.

## Acknowledgements

This work was funded by U.S. Food and Drug Administration Medical Countermeasures Initiative contract (75F40120C00085) awarded to JAH, ADD and DAM. The article reflects the views of the authors and does not represent the views or policies of the FDA. This work was also supported by the MRC (MR/W005611/1) G2P-UK: A national virology consortium to address phenotypic consequences of SARS-CoV-2 genomic variation (DAM, JPS, ADD and JAH), and received support from the European Union’s Horizon Europe research and innovation programme under grant agreement No 101057553 and the Swiss State Secretariat for Education, Re-search and Innovation (SERI) under contract number 22.00094 (AK). SG is funded by the China Scholarship Council/University of Bristol joint-funded PhD Scholarship programme. We are grateful to the technical staff in the Histology Laboratory, Institute of Veterinary Pathology, Vetsuisse Faculty, University of Zurich, for excellent technical support. We would also like to thank David Bauer and Harriet Mears (Francis Crick Institute, UK) for useful discussions on the RNA structure predictions.

## Figure legends

**Supplementary Figure 1.**
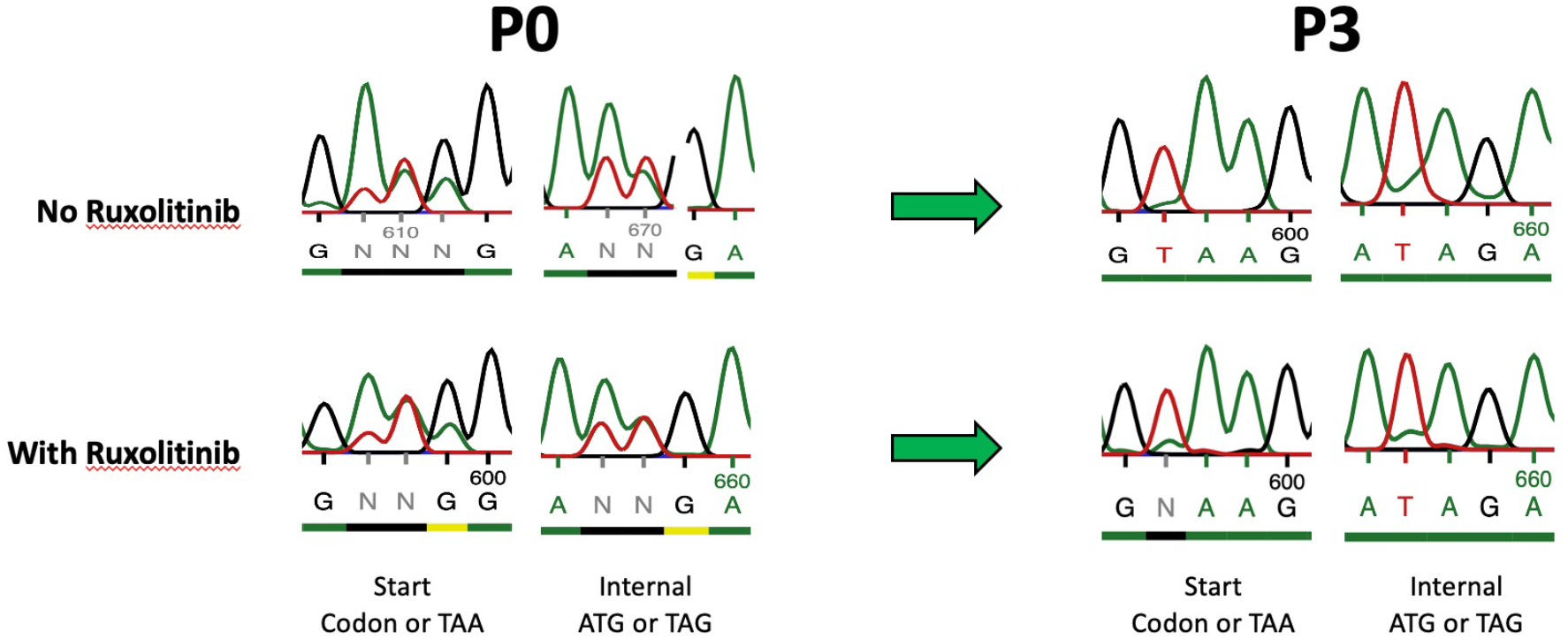
Effect of Ruxolitinib on competition assay. Sanger sequence data from a repeat of the competition assay in the presence or absence of the JAK/STAT pathway inhibitor Ruxolitinib. WT and ORF10KO virus were mixed in a 1:1 ratio and used to infect A549-AT cells previously treated with ruxolitinibor a mock carrier at an moi of 0.01. After 48 hours the supernatant virus was harvested, titrated and the infections (with or without Ruxolitinib as appropriate) repeated two more times. Samples of the P0 and P3 virus mixtures were sent for Sanger sequencing and the sequence data for the initiating and internal methionine codons are shown here.

## Tables

Supplementary Table 1. Quantitative proteomics analysis of A549-AT cells infected with WT or ORF10KO virus.

This analysis focuses on comparing ORF10KO and WT infected cells and shows log2 fold changes in abundance for human and viral proteins. In addition, normalised phospho-proteomic data is provided to quantitate relative changes in phosphorylation. The changes in phosphorylation are normalised relative to changes in protein abundance.

Supplementary Table 2. Quantitative list of all mutations observed within the ORF10 region in the whole genome amplicon sequence data from lungs of hamsters infected with WT or ORF10KO virus.

This analysis covers sequence data from whole genome Midnight amplicon sequencing. Within the spreadsheet there are 12 sheets that contain the analysis outputs for the six animals infected with the WT virus (WT1 – WT6) and for the six animals infected with the ORF10KO virus. On each sheet the sequences observed at the region covering the initiating codons are listed, a region that would be complimentary to the initiating codon and the region covering the internal AUG and its predicted stem loop. Each line of data indicates sequence data that is linked and came from the same sequence read and how often that pattern was observed.

Supplementary Table 3. Quantitative list of all mutations observed within the ORF10 region amplified by RT-PCR from lungs of hamsters infected with WT or ORF10KO virus.

This analysis covers sequence data from a RT-PCR product from just the ORF10 region. Within the spreadsheet there are 12 sheets that contain the analysis outputs for the six animals infected with the WT virus (WT1 – WT6) and for the six animals infected with the ORF10KO virus. On each sheet the sequences observed at the region covering the initiating codons are listed, a region that would be complimentary to the initiating codon and the region covering the internal AUG and its predicted stem loop. Each line of data indicates sequence data that is linked and came from the same sequence read and how often that pattern was observed.

